# A reinforcement learning based software simulator for motor brain-computer interfaces

**DOI:** 10.1101/2024.11.25.625180

**Authors:** Ken-Fu Liang, Jonathan C. Kao

**Affiliations:** Dept of Electrical and Computer Engineering, University of California, Los Angeles, CA, 90024, United States; Neurosciences Program, University of California, Los Angeles, CA, 90024, United States

## Abstract

Intracortical motor brain-computer interfaces (BCIs) are expensive and time-consuming to design because accurate evaluation traditionally requires real-time experiments. In a BCI system, a user interacts with an imperfect decoder and continuously changes motor commands in response to unexpected decoded movements. This “closed-loop” nature of BCI leads to emergent interactions between the user and decoder that are challenging to model. The gold standard for BCI evaluation is therefore real-time experiments, which significantly limits the speed and community of BCI research. We present a new BCI simulator that enables researchers to accurately and quickly design BCIs for cursor control entirely in software. Our simulator replaces the BCI user with a deep reinforcement learning (RL) agent that interacts with a simulated BCI system and learns to optimally control it. We demonstrate that our simulator is accurate and versatile, reproducing the published results of three distinct types of BCI decoders: (1) a state-of-the-art linear decoder (FIT-KF), (2) a “two-stage” BCI decoder requiring closed-loop decoder adaptation (ReFIT-KF), and (3) a nonlinear recurrent neural network decoder (FORCE).

## 1 Introduction

In intracortical motor brain-computer interfaces (BCIs), neural activity reflecting motor commands are decoded to control the movements of an end effector, such as a computer cursor on a screen or a robotic arm. Motor BCIs were initially demonstrated in the late 1990 and early 2000s^1–3^, entering pilot clinical trials in 2004^4^. The past two decades have seen significant advances to increase performance in movement decoding incorporating ideas from statistical inference^5–8^, feedback control^9–11^, dynamical systems^12,13^, closed-loop decoder and neural adaptation^14–16^, and deep learning^17–19^. However, BCIs remain in pilot clinical trials today, reflecting the relatively slow pace of BCI research where new innovations typically require months of experimental validation. Moreover, intracortical motor BCI research today is largely limited to a handful of labs with neurosurgically implanted subjects, further limiting the speed and community of BCI research.

A key reason motor BCI development takes months is that motor control tasks are fundamentally “closed-loop” (Figure 1a). The performance of a BCI relies on the user’s “control policy” used to interact with an imperfect BCI decoder. The user must constantly adjust his or her motor commands in response to feedback of the decoded output. This constant updating results in emergent interactions and unique control policies for each decoder, meaning decoder performance depends on these interactions. We emphasize that these control policies are also decoder dependent. For example, Figure 1b illustrates BCI experiments where monkeys were allowed to move their hands to control a velocity Kalman Filter (VKF)^20^ and a Feedback Intention Trained Kalman Filter (FIT-KF)^10^. The monkey’s ability to make overt hand movements was intentionally allowed in this experiment because these movements reflect motor commands used to control the decoder. The hand kinematics are very different for controlling the VKF versus the FIT-KF, with VKF exhibiting longer hand trajectories with erratic movements compared to FIT-KF (Figure 1c, see also Supplementary Video 1, FIT-KF average hand trajectory length: 202 mm, VKF: 468 mm, *p <* 10^*−*7^, Wilcoxon rank-sum test). Correctly predicting BCI performance therefore requires modeling how the user will uniquely interact with a particular decoder in closed-loop experiments.

**Figure 1.**
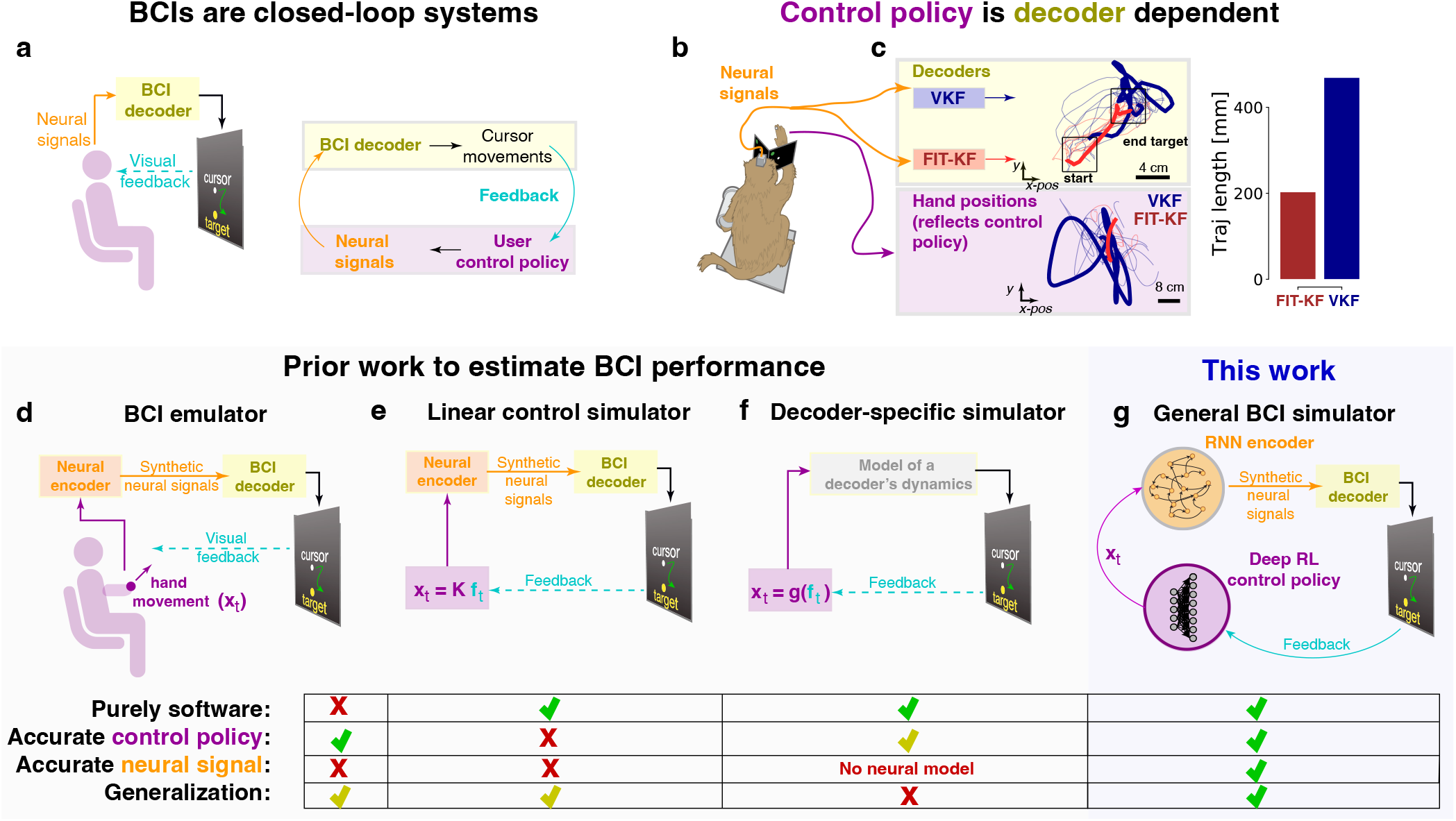
BCIs are closed-loop systems and prior work. **(a)** BCI decoders are imperfect, so decoded cursor movements will not fully match the intent of the user. In response, the user will generate updated motor commands and neural signals. **(b)** BCI performance relies on a decoder dependent control policy. Hand and cursor positions were recorded while the monkey neurally controlled a VKF and FIT-KF. **(c)** In the yellow box, we plot the decoded output of the VKF (blue) and FIT-KF (red) from intracortical BCI experiments where the monkey sought to acquire a target at 45^*?*^ (example trial bolded). In the purple box, we plot the monkey’s recorded hand positions, which reflect his motor commands to control the decoder. Hand trajectories when controlling the VKF range all over the workspace and have longer trajectories with erratic movements compared to FIT-KF. **(d)** The BCI emulator circumvents invasive neurosurgery by generating synthetic neural signals with hand kinematics, although it still requires human experiments to implement an accurate control policy. As we later show, the synthetic neural activity model (neural encoder) used by Cunningham et al. ^23^ is not accurate enough to reproduce past decoder studies. **(e)** The linear control simulator replaces the human with a linear control policy. This makes the simulator entirely software, but the control policy is not accurate. **(f)** The simulator by Willett et al. ^25^ enables hyperparameter optimization of a single decoder by modeling its dynamics (gray box), but cannot generalize to new decoders. **(g)** Our goal is the general solution, a purely software simulator that accurately predicts the performance of any type of decoder. **(d-f)** Green checkmarks indicate yes, red checkmarks indicate no, and yellow checkmarks indicate that while the answer is theoretically yes, in practice it is more nuanced and depends on a particular model.

Consistent with this, several studies document that analysis carried out in “offline” simulations that do not incorporate closed-loop feedback can be discordant with closed-loop experiments^18,21–24^.

While prior studies have attempted to emulate or simulate BCI systems, they face important limitations that we address. Cunningham et al. ^23^ developed a BCI “emulator” where a human subject’s hand movements generate synthetic neural activity to control a BCI decoder, removing invasive neurosurgery but still requiring experiments with a human-in-the-loop (Figure 1d, see Methods for more detail). We use the term “emulator” to highlight that this system requires a physical experiment with hardware that mimics the BCI decoding system and a human to provide a control policy. While this approach is useful, and we later use it to validate a generative model (neural encoder) which synthesizes neural activity, its use of human experiments limits its community use and speed. To remove the human-in-the-loop and avoid experiments, Lagang and Srinivasan ^26^ used the linear quadratic regulator (LQR) to approximate the human’s BCI control policy (Figure 1e). Other studies have also used linear policies from control theory in BCI design^27–31^. However, as shown by Willett and colleagues^25,32^, and by further experiments in this study, linear control policies are poor approximations of user control policies and result in incorrect conclusions. Finally, Willett et al. ^25^ designed a simulator that enabled hyperparameter tuning of a VKF (Figure 1f). This simulator, however, does not have a neural data model and requires closed-loop experiments to model a decoder’s dynamics. This simulator can therefore only optimize decoders already tested in closed-loop intracortical experiments and does not generalize to new, never-before-tested, decoders. It is also limited to optimize decoders of a linear form or a simple constrained nonlinearity.

Our BCI simulator aims to faithfully predict various BCI decoder performance entirely in software. We extend the ideas of Cunningham et al. ^23^ and Lagang and Srinivasan ^26^. Our simulator incorporates software models of synthetic neural activity and control policy (Figure 1g). In prior work, we showed that deep learning models could generate synthetic neural activity that models key features of motor cortical activity^33^. In this study, we improved these neural encoders to accurately replicate neural activity. This enhancement enables the BCI simulator to reproduce past closed-loop BCI studies. The critical and novel contribution of this work is to introduce a deep RL agent that replaces the need for a monkey-or human-in-the-loop, eliminating the necessity for physical experiments in our simulator. We observe that a more expressive deep RL agent is necessary, as simpler linear policies prove inadequate in controlling decoders, especially nonlinear ones. To our knowledge, this is the first demonstration of a versatile BCI simulator that uses both synthetic neural activity and an AI control policy to reproduce the results of published closed-loop BCI studies. Our simulator accurately predicts the detailed performance of several decoders, including those that require two stages of training, as well as linear and nonlinear decoders. Our simulator is implemented in software and can therefore be widely used.

## 2 Results

There are three components in a BCI: the decoder, neural activity, and user control policy. A general BCI simulator must accurately approximate the neural activity and control policy to correctly predict decoder performance. Our design approach uses (1) published BCI experiments, with *true* (not approximated) neural activity and *true* user control policy, (2) physical BCI emulator experiments, with *approximated* neural activity but *true* control policy, and finally (3) software BCI simulator experiments, with *approximated* neural activity and *approximated* control policy. Published BCI experiments provide ground truth decoder comparisons, collected with empirical neural recordings and a user-in-the-loop, that we use to validate our BCI emulator and simulator (Figure 2a). The BCI emulator, which approximates neural activity but incorporates a user-in-the-loop, allows us to validate the neural encoder in isolation to reproduce published ground truth decoder comparisons (Figure 2b,d). Finally, we build the BCI simulator by fixing a neural encoder from the BCI emulator and training deep RL agents to implement approximate control policies for each decoder, replacing the user-in-the-loop (Figure 2c,e). While constructing the simulator necessitates the initial use of an intracortical BCI dataset for a one-time training of a neural encoder, the entire BCI simulator is implemented purely in software and is accessible to everyone. Users of the future BCI simulator will not be required to conduct any physical experiments or have access to intracortical BCI data.

**Figure 2.**
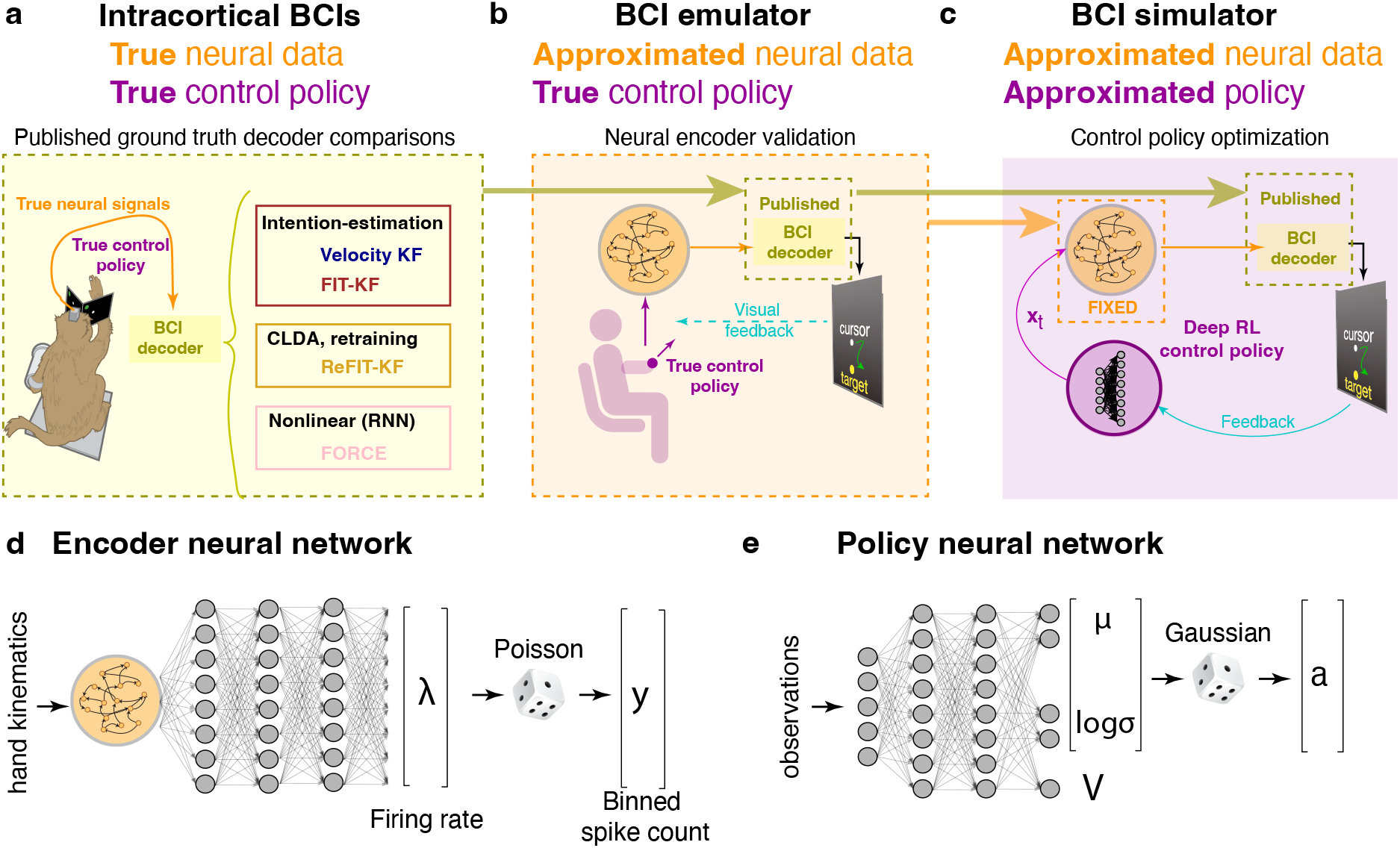
BCI simulator design approach. Our approach makes use of **(a)** published intracortical BCI literature, **(b)** BCI emulation, and **(c)** deep RL to design and validate the components of the BCI simulator. **(a)** Published BCI experiments, using true neural data and control policy, provide three different ground truth decoder comparisons to validate the BCI emulator and simulator. **(b)** The BCI emulator emulates monkey BCI experiments with true control policy from human testing subjects and synthetic neural activity from neural encoder. **(c)** We fix the neural encoder from the BCI emulator and then optimize the deep RL control policy in the simulator. **(d)** Architecture of the RNN neural encoder which takes hand kinematics and generates binned spike count. **(e)** Architecture of the deep RL policy which takes observations and generates hand acceleration and state value. Additional details for training RNN neural encoder and RL policy are provided in the Methods.

### 2.1 Ground truth decoder comparisons: true neural data, true control policy

We chose to replicate three distinct BCI studies that improved BCI decoders through different innovations. This was to demonstrate that our simulator, which did not incorporate any decoder-specific information, could make general and correct predictions of BCI decoder performance across diverse innovations and distinct study settings. We chose to replicate the BCI experiments that established the (1) FIT-KF^10^, (2) ReFIT-KF^9^, and (3) FORCE^17^ decoders. The FIT-KF is a linear state-of-the-art decoder whose high performance is due to the incorporation of control theory inspired dataset augmentation (also known as intention estimation)^10^. The ReFIT-KF, which also incorporates intention estimation, is a decoder that employs two stages of BCI training, a form of closed-loop decoder adaptation (CLDA)^15,34^. The ReFIT-KF decoder first requires the user to control a position velocity Kalman filter (PVKF), followed by a retraining stage using the neural activity and cursor movements during PVKF control^9^. Finally, the FORCE decoder is a nonlinear decoder that uses a recurrent echo state network (ESN)^17^. Our goal in choosing these studies was to demonstrate the simulator made correct predictions under very different decoder innovations, namely: (1) state-of-the-art linear decoding with intention estimation, (2) closed-loop decoder adaptation and retraining, and (3) nonlinear decoding. Additional decoder details are provided in the Methods.

### 2.2 Neural encoder validation: approximate neural data, true control policy

To validate the neural encoder, which maps motor commands (recorded hand kinematics) to synthetic neural activity, we employed the BCI emulator introduced by Cunningham et al. ^23^. The BCI emulator uses closed-loop experiments (Figure 2b). In this setup, a user moves his or her hand, generating synthetic activity that is subsequently decoded. Since the user observes the decoded output and adjusts his or her motor commands to better control the decoder, the BCI emulator incorporates a true (not approximated) user control policy. This setup allows us to evaluate the neural encoder in isolation. Further details of the BCI emulator are provided in the Methods. All experiments were approved by the UCLA IRB.

The prior study by Cunningham and colleagues used a Poisson process velocity tuning (PPVT) model of neural activity (see Methods), which they found resulted in successful optimization of decoder bin width. However, a PPVT neural encoder fails to capture the rich heterogeneity and dynamics of motor cortical activity^33^. We performed BCI emulator experiments using the PPVT neural encoder, and found it failed to reproduce published studies (Supplementary Figure 1), necessitating a better neural encoder.

#### 2.2.1 Improving the RNN neural encoder to more accurately replicate neural properties

While it may seem challenging to generate synthetic neural activity for successful use in BCI simulation, the properties of motor cortical activity simplify the problem. Specifically, motor cortical population activity is relatively low-dimensional^35–37^, exhibits structured dynamics^12,38,39^, and can reasonably be modeled by RNNs in tasks with simple inputs^40–43^. We hypothesized that a neural encoder that reproduces the heterogeneity, low-dimensionality, and dynamics of motor cortical neural populations during reaching would be capable of reproducing the three published decoder studies.

We trained RNN neural encoders to transform hand kinematic inputs to binned spike outputs (Figure 2d) using data collected from two Utah arrays while a monkey performed center-out-and-back reaches with their hand (details in Methods). Consistent with our prior offline work, we found these encoders better reproduced single electrode PSTHs, neural population trajectories, and neural population rotational dynamics in point-to-point reaches (Supplementary Figure 2a-c, Supplementary Table 1). RNN neural encoders also reproduced neural activity that could be accurately decoded offline, in contrast to PPVT (Supplementary Figures 3-4, Supplementary Tables 2-3). We additionally identified two important design considerations. First, it was important to incorporate a 200 ms delay between kinematics and neural activity, accounting for neural signal transduction from cortex to the muscles (Supplementary Figure 5). Second, it was important to regularize RNN input weights. This reduced the contribution of external inputs to RNN activity in comparison to internal RNN dynamics (Supplementary Figure 6), likely improving both neural population dynamics inference (Supplementary Figure 2d-i) and trajectory decoding (Supplementary Figure 4). Together, RNN-based neural encoders reproduced key features of motor cortical neural population activity, providing a better approximation of neural activity than the PPVT. We emphasize that the neural encoder is decoder-independent because it was trained with neural data and hand trajectories recorded in hand control experiments, as opposed to online BCI control experiments, which are decoder-dependent. This enables us to make a general neural encoder that would work well across decoder settings.

**Table 1.**
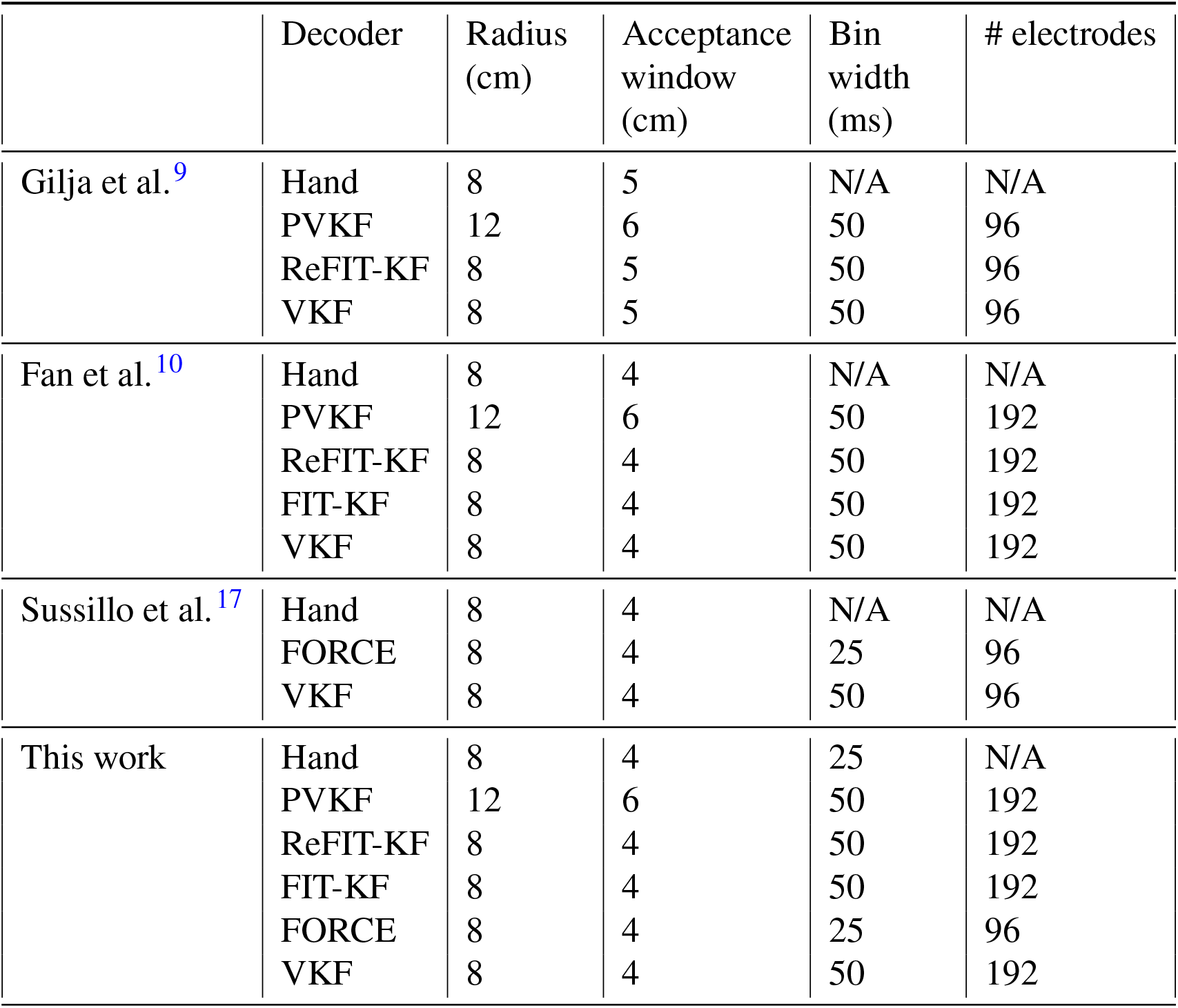
Task and recording settings for each of the three studies and this work. When the number of electrodes is 96, this corresponds to one Utah Array in primary motor cortex. 192 electrodes corresponds to two Utah Arrays, one in primary motor cortex and the other in dorsal premotor cortex.

**Figure 3.**
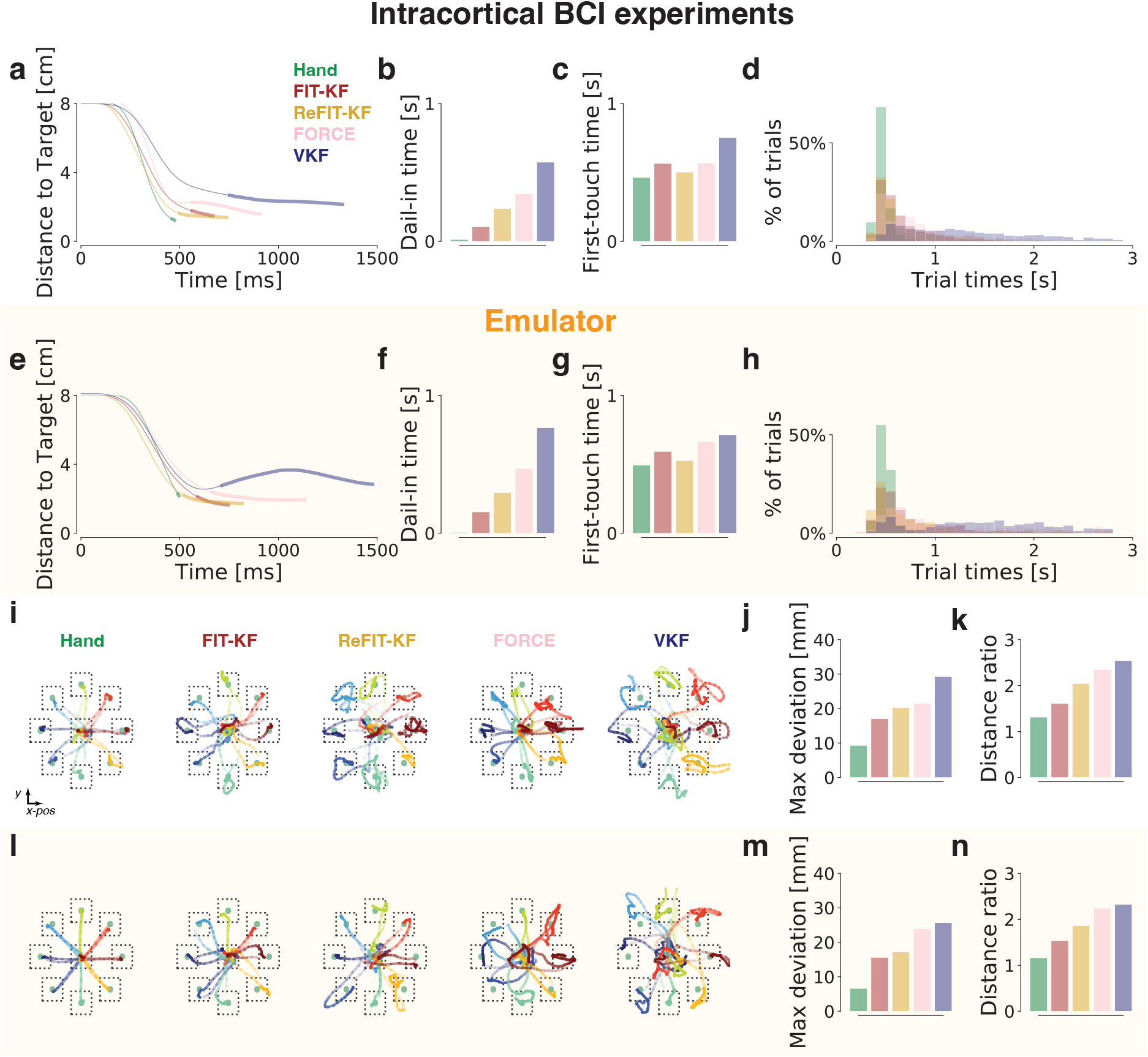
BCI emulator reproduces published studies. **(a)** Distance-to-target plots from intracortical BCI experiments for the various decoders. The bolded line corresponds to DIT. **(b, c)** Average DIT and FTT, respectively, from BCI experiments. **(d)** Distribution of trial times for each decoder. **(e-h)** Same as **(a-d)** but for the BCI emulator. **(i)** Randomly sampled trajectories for each decoder. **(j)** Max deviation from the straight line path for each decoder. **(k)** Distance ratio (cursor path length divided by straight path length) for each decoder. Distance ratio is the inverse of path efficiency. **(l-n)** Same as **(i-k)** but for the BCI emulator. Emulator results were shifted 200 ms to remove visuomotor response time when using the neural encoder. BCI and emulator results are summarized in Supplementary Table 4 and 5, respectively.

**Figure 4.**
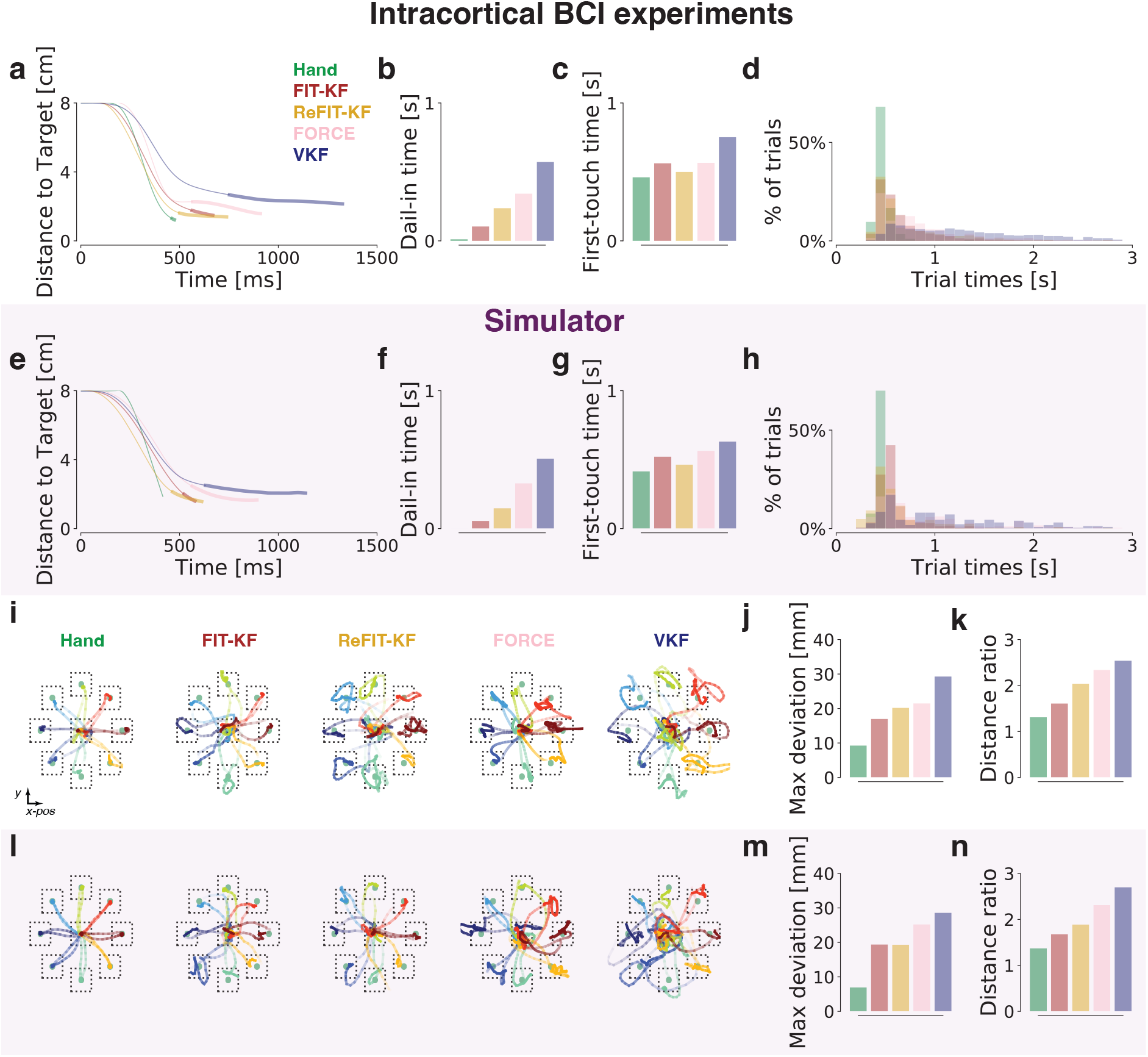
BCI simulator reproduces published studies. **(a-d, i-k)** Reproduction of the same panels from Figure 3, for ease of comparison of the BCI simulator to intracortical BCI experiments. **(e-h)** Same as **(a-d)** but for the BCI simulator. **(l-n)** Same as **(i-k)** but for the BCI simulator. BCI and simulator results are summarized in Supplementary Table 4 and 7, respectively.

#### 2.2.2 A BCI emulator using an RNN neural encoder reproduces published BCI studies

We performed the FIT-KF, ReFIT-KF, and FORCE studies in a BCI emulator using 4 human subjects. For each study, we reproduced the task, task parameters, decoder hyperparameters, and decoder training as described in the published studies^9,10,17^. We trained decoders as in the published studies by having subjects first perform center-out or pinball task reaches, followed by supervised training with the concurrent kinematics and synthetic neural activity generated by the neural encoder (see Methods). The results of the published studies are combined in Figure 3a-d, i-k. Each study reported superior performance over the then state-of-the-art VKF, due to superior fine control to successfully hold the cursor over the target (Dial-In Time or DIT, Figure 3a,b, see Supplementary Figure 7a for further description) and faster time to first touch the target (First Touch Time or FTT, Figure 3a,c, see Supplementary Figure 7b for further description). This also led to smoother kinematic trajectories, with improved path efficiency and less deviation from the straight-line path (Figures 3i-k, see Methods).

Our BCI emulator, using an RNN neural encoder, reproduced the key conclusions of each study. Surprisingly, it also produced detailed timing and trajectory results across all studies. The summary of our decoder comparisons are in Figure 3e, where we reproduced the distance-to-target plot results reported in all studies. This is also shown in Supplementary Video 2. In BCI emulator experiments, we highlight the following: (1) we reproduced that FIT-KF, ReFIT-KF, and FORCE all achieved superior performance to the VKF, as well as the result in Fan et al. ^10^ that FIT-KF and ReFIT-KF achieve similar trial times with ReFIT-KF having faster FTT than FIT-KF, but FIT-KF having better DIT than ReFIT-KF (Figure 3a,e); (2) we reproduced the ordering of fine control DIT across studies, in particular that FIT-KF had the shortest DIT, followed by ReFIT-KF, FORCE, and VKF (Figure 3b,f); (3) we reproduced the trends in FTT across studies, where VKF had the slowest FTT, ReFIT-KF had the best FTT, and FIT-KF and FORCE were intermediate (Figure 3c,g); (4) we achieved similar trial time distributions over single-trials across all decoders, with the VKF having the widest trial-time distribution (Figure 3d,h and Supplementary Figure 8 for one-by-one comparison); (5) we observed qualitatively similar decoder trajectories (Figure 3i, l) and reproduced the decoder ordering of max deviation and distance ratio (Figure 3j,k,m,n). Together, these results show that our BCI emulator with an RNN neural encoder reproduced the key timing and kinematic performance statistics of decoders evaluated in intracortical experiments, including timing at the single-trial resolution.

We emphasize that the neural encoder was only trained from open-loop center-out-and-back reaches. Remarkably, when used in BCI emulation for very different studies, we observed this approximate neural data was sufficient for predicting BCI decoder performance. We emphasize that these studies vary significantly; for example, ReFIT-KF emulation required an intermediate stage of decoding with a PVKF using the neural encoder, followed by subsequent retraining, whereas the FORCE and FIT-KF were nonlinear and linear decoders, respectively, trained directly from hand movements. We found that the intermediate PVKF training stage of the ReFIT-KF also closely matched PVKF performance from intracortical experiments (Supplementary Figure 9 and Supplementary Video 2). By closely modeling the kinematic-to-neural relationship, we were able to accurately reproduce detailed timing and kinematic performance of these decoders in BCI emulation, in contrast to a poorer PPVT model. These results suggest that an RNN neural encoder is sufficient to accurately predict decoder behavior for 2D cursor control.

### 2.3 Replacing the user-in-the-loop: Approximate neural data, approximate control policy

We next sought to replace the user-in-the-loop in BCI emulation with an AI agent. The AI agent must interact with the BCI system to acquire targets, as humans did in the BCI emulator. Because the BCI emulator reproduced published studies, we reasoned that successful replication of user control policy would lead to a purely software BCI simulator that also reproduces published studies. We therefore fixed the neural encoder validated in BCI emulation, enabling sole optimization of an agent.

There are important considerations for the design of such an agent. First, as we demonstrated in Figure 1b, control policy is *decoder dependent*. The agent architecture must therefore be expressive enough to be able to control a variety of decoders. Second, the control policy should be able to control decoders like monkeys and humans do. The agent should therefore be subject to constraints, for example, that it does not make non-physiological accelerations.

We first considered whether a linear quadratic regulator (LQR) was sufficient to reproduce the results. However, we found that LQR failed to control all decoders, which is not entirely surprising because the overall system is nonlinear. This is consistent with prior work that showed linear control policies are inadequate for BCI control^25^. When we relaxed the neural encoder to a PPVT, an LQR agent was able to control it. However, an LQR agent with a PPVT neural encoder results in discordant conclusions compared to prior studies (Supplementary Figure 10).

We therefore trained nonlinear control policies implemented via deep neural networks. The AI agent must generate movements resulting in similar online decoder performance as in intracortical experiments. While imitating human or monkey control policies with empirical online control data may be helpful in reproducing published studies, imitation learning would limit the simulator’s ability to generalize to never-before-tested decoders, where online control data is not available. The agent had to learn to control each decoder anew, some decoders of which may have never before been tested. We therefore trained a deep RL agent, where the reward (successfully acquiring a target) was sufficient to guide training of new decoder-dependent control policies. We trained using proximal policy optimization (PPO), an actor critic method (Figure 2e), to maximize expected reward.

Out-of-the-box trained PPO agents did not successfully replicate experimental results because naïve deep RL agents critically do not have any biophysical constraints on their actions. They can generate arbitrary acceleration and deceleration, and further do not have any constraints to reduce unnecessary movements. As such, we found an unconstrained deep RL agent did not reproduce prior BCI studies (Supplementary Figure 11). To address this problem, we therefore regularized training by introducing two new KL divergence constraints that encouraged the RL agents to have smooth actions and minimize energy use (see Methods). These constrained the agent to generate more plausible accelerations and not move unnecessarily. Finally, we also found it was important to use curriculum learning to facilitate training deep RL agents to reproduce the prior studies. Further details of these regularizations and training, as well as a more detailed description of the RL problem and hyperparameters, are in the Methods.

#### 2.3.1 A software BCI simulator with a deep RL agent reproduces published BCI studies

We trained the deep RL agent to control each of the decoders in the published studies. Because control policy is decoder dependent, the deep RL agent was trained to learn the emergent closed-loop interactions helpful for controlling each decoder. We found that the deep RL agent exhibited qualitatively similar decoder dependent control policies to monkeys controlling intracortical BCIs and humans controlling the BCI emulator (Supplementary Video 3, FIT-KF average “hand” trajectory length: 284 mm, VKF: 373 mm, *p <* 10^*−*7^, Wilcoxon rank-sum test).

We performed the FIT-KF, ReFIT-KF, and FORCE studies in a BCI simulator using trained RL agents.

We found the BCI simulator reproduced all the key conclusions of these studies, like in the emulator (Figure 4 and Supplementary Figure 9 for PVKF), including detailed timing and kinematic differences in decoder comparisons. An example is shown in Supplementary Video 4. This included correctly predicting the trends in first-touch time and dial-in time across all decoders, as well as kinematic trends in distance ratio and max deviation. Together, these results show the BCI simulator, using a deep RL agent, and an RNN neural encoder, reproduced the key timing and kinematic performance statistics of decoders, including timing at the single-trial resolution. This validates that our BCI simulator – implemented entirely in software – arrived at the same conclusions as prior intracortical studies without requiring months to years of invasive experiments.

## 3 Discussion

We incorporated a deep-learning-based neural encoder and deep RL agent to build a BCI simulator entirely in software without requiring invasive neurosurgery or human-in-the-loop experiments. Our critical innovation was replacing the BCI user with a deep RL agent trained with biophysically-inspired constraints to implement a purely software simulator. This simulator reproduced three prior BCI studies with distinct decoders using a general neural encoder trained with decoder-independent neural data.

Our work significantly extends prior attempts at BCI simulation and emulation. Our new RNN neural encoder better recapitulates important neural features which can be faithfully decoded to simulate cursor movement dynamics than prior works. PPVT neural encoder cannot provide critical neural features such as neural dynamics. Prior RNN neural encoder cannot generate precise neural activity and our innovations are necessary. We emphasize that it was important to incorporate the delay between hand speed and neural activity in the neural encoder, as well as regularize the encoder input weights (Supplementary Figures 2-6). The neural encoder can be built once from intracortical recordings of a subject performing a center-out-and-back task (or any task with simultaneous kinematics and neural activity). Our neural encoder was not trained with any decoder-specific knowledge, therefore, our work does not require new intracortical experiments to test new decoders: the fixed neural encoder built from point-to-point reaches serve as a proxy of a monkey brain can reproduce the same conclusions across 3 distinct BCI studies.

Our BCI emulator demonstrated that true control policy coming from human testing subject with RNN neural encoder can reproduce BCI studies, however, naive LQR used in Lagang and Srinivasan ^26^ cannot control the nonlinear BCI systems and better control policies are needed. Model-based control policies can be applied on applications that can be modeled as Markov decision processes (MDPs) including Backgammon, Go, robot locomotion, helicopter flight, autonomous driving, etc. MDP models may be computationally infeasible to use directly, but can be estimated through sampling. Model-free controlpolicy is more general and has been shown capable of learning human motor policies and skills in various tasks^44–48^. We used state-of-the-art RL algorithm, PPO, as an example to show the feasibility of building BCI simulator. We demonstrated that, in BCI simulation, constrained deep RL agent can naturally control nonlinear BCI systems as in BCI emulation and reproduce decoder performance. We emphasize the importance of the regularizations we introduced to enforce smoothness and low-energy actions from the agent; without these, the agent could make non-physical arbitrary accelerations and decelerations.

Our simulator accurately estimates the performance of linear and nonlinear decoders, in addition to those that require two training stages (closed-loop decoder adaptation). Our AI agent architecture, hyperparameter, and training approach were the same across all experiments, indicating that our BCI simulator generalized to different decoder algorithms. In contrast to time-consuming BCI experiments, our BCI simulator can benchmark decoder algorithms across different tasks, addressing a limitation where prior BCI literature tests different decoders, obfuscating comparison. For example, BCI simulator can evaluate the robustness of decoding algorithms by manipulating neural encoder to artificially simulate the neural recording instabilities such as baseline shift, unit drop-out, and tuning changes^49^. Baseline shifts were generated by applying a different random constant to the spike firing rate on each channel. Unit drop-out instabilities were generated by setting the activity of a subset of electrodes to 0. Instabilities resulting in tuning changes were generated by replacing the activity of a subset of electrodes with that of a held-out set of channels. Another approach to simulate the neural recording instabilities is to train multiple neural encoders from the same monkey data cross days. Each neural encoder serves as a proxy of a monkey brain on the day of data collection. Also, general decoder allows us to plug-and-play, however, it requires multiple testing subjects to evaluate such decoder. In our work, we can build multiple neural encoders from multiple subjects, and benchmark decoders across separate encoders in parallel simulation.

Online decoder performance depends on decoder algorithms and also other different perspectives including user-learning and neural plasticity. User-learning was considered by incorporating true human control policy in BCI emulator and training RL agents in BCI simulator. Our neural encoder serves as a proxy of the monkey brain that we recorded. The neural encoder was fixed in both BCI emulator and BCI simulator. Although it is possible to train multiple neural encoders with neural recordings from multiple days to simulate long-term neural variance. However, an important limitation of BCI emulator and simulator is decoder-dependent neural plasticity. Therefore, if a novel decoder algorithm incorporates neural plasticity or aspects of neural variance not captured by our neural encoder, then the simulator may not adequately evaluate this decoder’s performance. To further support large scope of decoding algorithms, new neural encoder architecture and neural plasticity mechanism may be needed in our future work.

This work also proves the concept that synthetic neural activity and deep RL agents can be combined to accurately simulate cursor control BCIs in 2D tasks. Future work may therefore apply our framework to other BCIs. The basic architecture of our simulate is generalizable to higher degree-of-freedom (DoF) BCIs, such as those that interact with a robotic arm or hand. The system requires an initial dataset of simultaneous recordings and the controlled output (e.g., multi-DOF kinematics or kinetics for a robotic arm) to do a one-time build of a neural encoder for the system. The amount of data required is not very large; in our system, our neural encoder was robust from training with only 500 total trials of center-out-and-back reaches, approximately 10 minutes of wall time experimental data. Following this, the RL agent must be trained to control the new BCI system with potential biomechanical models to constrain higher DOFs as demonstrated in Fischer et al. ^44^, Joos et al. ^45^, Cheema et al. ^50^.

## Acknowledgments

We thank Krishna V. Shenoy for intracortical BCI data, and Sergey Stavisky for helpful discussions regarding the manuscript. This work was supported by the following awards to JCK: National Institutes of Health R01NS121097 and DP2NS122037, and a UCLA Computational Medicine AWS grant. KL was supported by Study Abroad scholarship from ministry of education in Taiwan. We gratefully acknowledge the support of NVIDIA Corporation with the donation of the Titan Xp GPU used for this research.

## Author contributions statement

KL and JCK conceived of and designed the BCI simulator. KL performed BCI emulator experiments and optimized the neural encoder and control policy. KL and JCK wrote the manuscript. KL and JCK were involved in all aspects of the study.

## Data availability

All emulator and simulator data are available upon request.

## 4 Methods

Our Methods are organized as follows: (1) task descriptions, (2) dataset descriptions, (3) BCI decoder algorithms, (4) metric definitions, (5) BCI emulator description, (6) neural encoders, and (7) BCI simulator control policy and deep RL training.

### 4.1 Tasks

The three published studies we used to validate our emulator used variants of the center-out-and-back task and the pinball task. The center-out-and-back task was used to collect training data for PVKF, ReFIT-KF, FORCE, and VKF decoders. The pinball task was used to collect training data for FIT-KF decoder. All studies used the center-out-and-back tasks to quantify decoder performance. The center-out-and-back task parameters for each study and our work are summarized in Table 1.

#### 4.1.1 Center-out-and-back task

In the center-out-and-back task, eight “radial targets” were uniformly placed on the circumference of an 8-cm or 12-cm radius circle. One “center target” was placed at the center of the circle. Each target had a square acceptance window centered around the target, with side length 4, 5, or 6 cm depending on the study. The subject had to hold the cursor within the target acceptance window for 500 contiguous milliseconds to successfully acquire the target. After acquisition of the center target, one of the eight radial targets would be randomly prompted. The target had to be acquired within 3 seconds, or the trial was counted as a failure. After a successful acquisition of a radial target, or following the failure to acquire any radial target, the center target was prompted.

#### 4.1.2 Pinball task

In the pinball task implemented by Fan et al. ^10^, targets were randomly prompted in a 20-by-20 cm workspace. The target’s position was randomly sampled in each trial. Each target had a 4 cm square acceptance window. The subject had to hold the cursor within the target acceptance window for 750 contiguous milliseconds to successfully acquire the target. The target had to be acquired within 3 seconds, or the trial was counted as a failure. After a successful acquisition or following the failure to acquire any target, the next random target was prompted. There were two additional constraints on the next target location. First, the minimum distance between each random target was 4 cm to avoid two successive trials having overlapping acceptance windows. If the next target was within 4 cm of the previous target, the next target position would be re-sampled until the next target was more than 4 cm away. Second, if the next target was more than 14 cm away from the previous target, the next target position would be re-sampled until the next target was less than 14 cm away.

### 4.2 Dataset descriptions

We validated our simulator using results from the published studies of Gilja et al. ^9^, Fan et al. ^10^, Sussillo et al. ^17^. In these intracortical experiments, monkeys were trained to control BCIs to perform the center-out-and-back task. We used concurrent kinematics and intracortical threshold crossings recorded when Monkey J performed center-out-and-back task with his own hand as training dataset to train neural encoder. We also used the same day recorded online BCI data where Monkey J used FIT-KF and ReFIT-KF to perform the task as validation dataset for evaluating generalization performance of neural encoders. Finally, we merged the BCI data from Fan et al. ^10^ (Hand, FIT-KF, ReFIT-KF, PVKF, VKF) and Sussillo et al. ^17^ (FORCE) to obtain ground truth decoder performance.

In these studies, Monkey J had two Utah arrays, one in M1 and the other in the dorsal aspects of the premotor cortex (PMd). The recorded neural activity were threshold crossings, with the per-electrode threshold set at *−*4.5*×* the root mean square voltage of the electrode. For BCI decoders, the number of electrodes were either 96 (M1) or 192 (M1 and PMd), and spike counts were binned in either 25 or 50 ms bins depending on the study (see Table 1). In each of our experiments, we matched the number of electrodes used by the BCI experiments that we compare. FIT-KF, ReFIT-KF, PVKF, and VKF were using 192 electrodes, and FORCE was using 96 electrodes.

During hand and BCI control in these experiments, hand position was recorded whenever the monkey’s hand position was in view of an IR detector at a rate of 60 Hz (Polaris tracking system, NDI, Ontario, Canada). Hand position during hand control, with concurrent threshold crossings, comprised supervised training data that were used to train the neural encoder. The analyses that control policy is decoder-dependent in Figure 1 used recorded hand positions during VKF and FIT-KF control.

### 4.3 BCI decoder algorithms

Because each BCI decoder algorithm has been well-documented in its published studies, we comment on these more briefly in our Methods, summarizing their innovations and highlighting key differences between the decoders. The FIT-KF, ReFIT-KF, and VKF are all based on a linear dynamical system model where the cursor kinematics at time *t*, **x**_*t*_, are the state and binned spike counts at time *t*, **y**_*t*_, are the observations. **y**_*t*_ was a 96D or 192D vector containing the binned spike counts of 96 or 192 electrodes at time *t* (depending on the study), and **x**_*t*_ was a 5D vector representing the cursor’s x-and y-positions, velocities, and a bias term of 1 to model a baseline spike count. The dynamical system is:

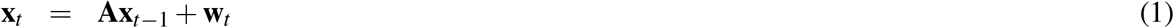

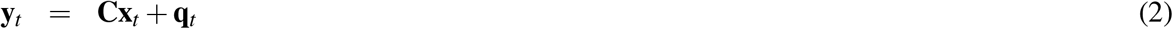

where **w**_*t*_ *∼ 𝒩* (**0, W**) and **q**_*t*_ *∼ 𝒩* (**0, Q**) are Gaussian noise. The parameters of this model, *θ* = *{***A, C, W, Q***}*, were inferred using supervised learning with maximum-likelihood estimation, where datasets comprised the paired cursor kinematics and neural activity, *{***x**_*t*_, *y*_*t*_*}*_*t*=1,…,*T*_. For more details, refer to Wu et al. ^5^, Gilja et al. ^9^, Kim et al. ^20^, Kao et al. ^51^.

To decode, we used the Kalman filter algorithm to infer the state of the dynamical system, 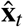, given the observed spike counts **y**_*t*_. This enabled recursive estimation of the state, 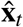, from a new neural observation, **y**_*t*_, and the previously decoded state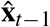. In the FIT-KF and ReFIT-KF, the Kalman filter recursion included a causal intervention, as well as a position correction, detailed further in Gilja et al. ^9^. The VKF did not include these.

#### 4.3.1 VKF and PVKF

The Velocity Kalman filter (VKF) and Position Velocity Kalman filter (PVKF) are linear decoders trained from concurrent cursor kinematics and spiking activity. Across all studies, and in our emulator and simulator, the VKF and PVKF were trained by collecting approximately 500 trials of a center-out-and-back task where targets were on a 12-cm radius circle with 4 cm square acceptance windows. VKF and PVKF in all studies used spike counts binned in non-overlapping 50 ms bins. The PVKF decoder models neural activity as a function of both position and velocity while the VKF only models it as a function of velocity. The VKF and PVKF do not incorporate intention estimation or closed-loop decoder adaptation^9,10^.

#### 4.3.2 FIT-KF

The Feedback Intention Trained Kalman filter (FIT-KF) is a linear decoder trained from concurrent cursor kinematics and spiking activity. The FIT-KF is trained from approximately 500 trials of the pinball task. FIT-KF training includes intention estimation, a form of dataset augmentation where cursor velocities in the training set are rotated to point towards the target at every time step. When the cursor is in the target acceptance window, intention estimation sets the cursor velocity to zero. These changes assume that at every point in time, the BCI user is giving motor commands to move the cursor towards the target, and that when in the acceptance window, the BCI user is attempting to hold the cursor still over the target. The FIT-KF also pre-processes the data to exclude the first 250 ms of data at the start of a trial, since this corresponds to the reaction time of the monkey. It also excludes the dial-in time (between when the user first touched and last touched the target) from training. The FIT-KF inference used a causal intervention in Kalman filter inference, where the covariance of the state position estimate is set to zero, since the user sees the true position of the cursor. Finally, the FIT-KF subtracts the contribution of position to the neural activity through a linear mapping, the details of which are described in Fan et al. ^10^.

#### 4.3.2 ReFIT-KF

The Recalibrated Feedback Intention-Trained Kalman Filter (ReFIT-KF) is a linear decoder that involves two stages of training, a form of closed-loop decoder adaptation. Concurrent cursor kinematics and spiking activity were used to train a Position Velocity Kalman Filter (PVKF), using the same task as used to train the VKF. Subsequently, the PVKF was used in closed-loop control to collect 500 trials of a center-out-and-back task where the target radius was 12 cm and each target had a 6 cm square acceptance window. The ReFIT-KF was then trained from the closed-loop 500 PVKF center-out-and-back trials. The PVKF data was augmented to include intention estimation. The ReFIT-KF inference also used the causal intervention and position contribution subtraction in the FIT-KF, the details of which are described in Gilja et al. ^9^.

#### 4.3.4 FORCE

The FORCE decoder is a nonlinear decoder. The FORCE decoder is a recurrent neural network (RNN) that takes the form of an echo state network (ESN). The FORCE decoder was trained using concurrent cursor kinematics and spiking activity using the same task used to train the VKF. The parameters of the FORCE decoder were trained using FORCE learning^52^ and the network was trained on four passes through the data, the details of which are described in Sussillo et al. ^17^.

### 4.4 BCI metrics

We describe the following metrics used to quantify and compare encoder and decoder performance.

**Contribution ratio** is the ratio of contribution from external input versus internal recurrent dynamics to the RNN states. The ratio is defined as

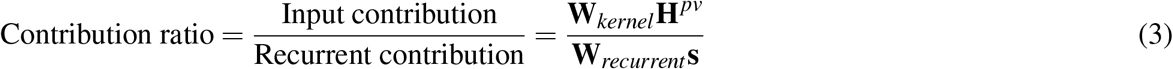

where **W**_*kernel*_ and **W**_*recurrent*_ are the input kernel (or input weight matrix) and recurrent weight matrix of the RNN, respectively. **H**^*pv*^ is the neural encoder kinematic input, which includes hand position and velocity, and **s** is the RNN internal state. We calculated contribution ratio for each artificial neuron and averaged across all neurons. A ratio close to zero means that the recurrent contribution dominates the state changes. In contrast, a large ratio means that the kinematic input dominates the state changes.

**Pearson correlation coefficient (PCC)** of PSTHs and neural trajectories: We report the PCC between BCI and synthetic PSTHs. The data were binned at intervals of 25 ms. For each electrode, we concatenated the PSTH for each of eight target reach conditions into a vector. We then calculated the PCC between these vectors for real and synthetic PSTHs, and average across electrodes. We also report the PCC between BCI and synthetic neural trajectories from PCA. PCA is an orthogonal transformation of the neural data that maximizes the variance of the data in low dimensional projections. We performed PCA on PSTHs to emphasize across-condition variability. When calculating PCC of neural trajectories, we projected both real and synthetic neural population activity into the PCs found from real data.

**Normalized root-mean-square error (NRMSE)** of PSTHs and neural trajectories: NRMSE is defined as

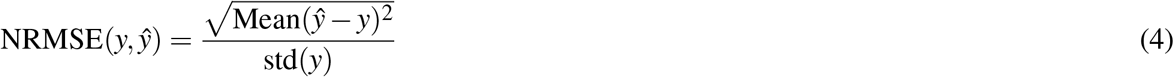

where *y* is the ground truth, ŷ is the predicted output, and “std” computes standard deviation. To compute the NRMSE of PSTHs, we first concatenated the PSTH for each of eight target reach conditions into a vector on a single electrode basis. We then calculated the overall NRMSE by averaging the NRMSE across all electrodes. For neural trajectories, we similarly projected both real and synthetic PSTHs into the PCs found from real data. We then calculated the overall NRMSE by averaging the NRMSEs across the top *N* dimensions. The value *N* is the number of PCs that capture over 90 % of the PSTH variance.

**jPCA** is a rotation of the top PCs that reveals rotational structure in data. We applied jPCA after finding top PCs. This comprised finding a skew symmetric matrix least-squares mapping from the position of the neural trajectory to its velocity, with additional details in Churchland et al. ^38^. We report 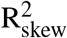, which is the variance explained in predicting the neural trajectory velocity from its position. A higher 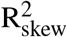 indicates that the system is better described by rotational dynamics.

**NRMSE of decoded velocity from real and synthetic neural activity** We quantify the NRMSE for decoding real and synthetic neural activity to evaluate the performance of a neural encoder. Neural activity from the monkey and neural encoder was decoded into cursor velocity. We then calculated NRMSE of cursor velocity for each trial and averaged across trials. NRMSE follows the Equation 4.

**Dial-in time**(DIT) is the time from when the cursor first enters the target acceptance window to when it last enters the target acceptance window prior to successfully holding the target. If the cursor enters the acceptance window and successfully holds the target without ever leaving the acceptance window, DIT is equal to zero. An illustration of the DIT is shown in Supplementary Figure 7a.

**First touch time**(FTT) is the time from trial start to when the cursor first enters the target acceptance window. An illustration of FTT is shown in Supplementary Figure 7b.

**Distance ratio**, also known as the inverse of path efficiency, is the distance traveled by the cursor divided by the straight line distance (i.e., the shortest trajectory) from the start point to the end point of a trial. Distance ratio quantifies path inefficiency. When the distance ratio is large, then the path taken to acquire the target is less direct.

**Maximum deviation** is the maximum distance between the cursor trajectory and the straight line from the start point to the end point of a trial. Maximum deviation therefore quantifies how far the cursor deviates from the straight line.

### 4.5 BCI emulator

We built a BCI emulator in a similar fashion to the “Online Prosthetic Simulator” introduced by Cunningham et al. ^23^. In our BCI emulator, endpoint kinematics of the user’s hand were recorded via the Polaris Vega system (Northern Digital, Ontario, Canada). This system tracked the position of an infrared bead to a resolution of 0.30 mm at 250 Hz, enabling precise estimation of hand position. These data were processed by a custom real-time system build using the LiCoRICE framework^53^, which used 1 ms clock cycles, enabling simulation of spikes-based BCIs. Our real-time system processed hand kinematics and transformed them into spike counts via the neural encoder. The synthetic neural spike counts were decoded to update the cursor positions shown on a screen. In response to this visual feedback, the user made new motor commands to best control the BCI.

In experiments, we had each subject perform approximately 50 native arm reaches to gain familiarity with the system. We then collected training data for 500 native arm reaches on center-out-and-back and pinball tasks, respectively. This data, comprising paired cursor kinematics and synthetic binned spike counts, were used to train the FIT-KF, FORCE, VKF and PVKF decoders. To train the ReFIT-KF, we collected 500 reaches in PVKF-controlled center-out-and-back task. Testing subjects practiced decoders in a sequence (arm, FIT, ReFIT, FORCE, then VKF) until they felt confident in controlling decoders. Following this, in evaluation sessions, subjects follow the same sequence (arm, FIT, ReFIT, FORCE, then VKF), controlling each decoder for 60 center-out-and-back reaches. Informed consent was obtained from all human research participants. Subjects were notified they could stop the experiment when they felt fatigue. We repeated this sequence until the subject terminated the experiment due to fatigue.

### 4.6 Neural encoders

The neural encoder transforms hand kinematics into binned spike rates. Neural encoders were trained on recorded data when monkey performed center-out-and-back task with his arm, and further evaluated with validation dataset recorded when monkey controlled FIT-KF and ReFIT. In total, we used approximately 500 successful trials to train and test neural encoders, and 300 (300) successful trials from FIT-KF (ReFIT-KF) experiments as validation dataset.

#### 4.6.1 PPVT neural encoder

The PPVT models neural firing rates based on a tuning curve model and reach speed. The tuning curve quantifies each neuron’s firing rate as a function of reach angle, with the angle eliciting the highest firing rate called the “preferred direction” (PD). Firing rate is then linearly scaled based on the speed of reach.

In total, the firing rate is calculated as

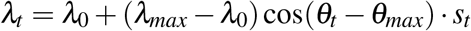

where *λ*_0_ is the neuron baseline firing rate, *λ*_*max*_ is the neuron’s maximum firing rate, *θ*_*t*_ is the reach angle at time *t, θ*_*max*_ is the neuron’s preferred direction, and *s*_*t*_ is proportional to the reach speed at time *t*. We additionally scaled *s*_*t*_ so that firing rates, when decoded, produced reasonable trajectories. This scaling is subject dependent, since different subjects may reach with different vigor. After computing firing c to an inhomogeneous Poisson Process, as below:

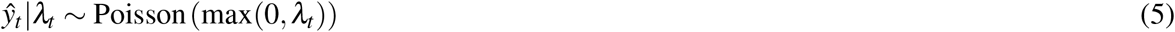

where *ŷ*_*t*_ is a vector of the binned spike counts for each neuron at time *t*.

#### 4.6.2 RNN neural encoder

Our RNN model was trained in a supervised manner with recorded hand kinematics comprising 2D position and velocity (inputs) and binned spike counts (outputs) in the monkey center-out-and-back task with arm reaches. Kinematics and binned spike counts were evaluated at 25 ms bin width resolution. The overall neural encoder used one RNN layer followed by three multilayer perceptron (MLP) layers. We use superscript *f* and subscript *N* to denote activation function, *f*, and number of neurons, *N*, for each layer, respectively. We used the sigmoid activation function, i.e., 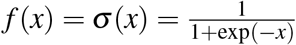 in all hidden layers. Because firing rates are non-negative and not bounded by 1, we used an exponential nonlinear activation function in the final layer, i.e., *f* (*x*) = *exp*(*x*). The complete RNN neural encoder is

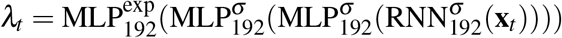

with spikes generated according to equation 5. The RNN was trained in “seq2seq” fashion by maximizing the log-likelihood of the observed binned spike counts under the assumption that the spike counts are Poisson distributed.

As described in the paper, we also regularized the input weights (also known as kernel weights) of the RNN, which we refer to as the “regularized RNN neural encoder.” In addition, instead of using concurrent hand kinematics and binned spike counts to train neural encoder, we also delayed binned spike counts for 200 ms, which achieved the highest correlation coefficient to hand speed (Supplementary Figure 5). This regularized RNN neural encoder trained with delay binned spike counts is called the “delayed regularized RNN neural encoder” and was used in both simulator and emulator. All RNN neural encoders were optimized with stochastic gradient descent using the Adam optimizer, and our network was initialized with the Xavier uniform initialization. Gradients for stochastic gradient descsent were computed using backpropagation-through-time.

### 4.7 Simulator

The simulator replaces the user-in-the-loop with a deep RL agent. It therefore allows us to predict the performance of BCIs without human subjects. In this setting, the hand kinematics (input of the neural encoder) are controlled by the deep RL agent, instead of a human. The rest of the simulator mimics the BCI emulator. The hand position, 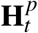, and velocities, 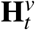, of the agent are updated according to the following kinematic equation:

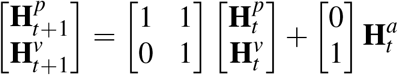

where 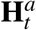 is the hand acceleration at time *t*. The synthetic spike counts were synthesized with hand kinematics and decoded by decoders into cursor position as the following:

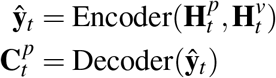

Our goal was to therefore have the agent generate hand accelerations, 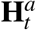, that mimicked how a user would control a BCI. The input to the agent, also called the “observation,” was the hand’s position and velocity, cursor’s position and velocity, as well as the target position. The output of the agent was a hand acceleration that would subsequently be translated into neural activity, **ŷ**_*t*_, and decoded into a new cursor position,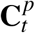.

#### 4.7.1 LQR

The linear quadratic regulator (LQR) algorithm is an automated way of finding a state-feedback controller based on system dynamics and cost constraints. System dynamics are described by a set of linear differential equations and cost constraints are described by quadratic functions. In our tasks, the observation space was the concatenation of cursor position, cursor velocity and target position. The action space was the hand acceleration.

The LQR assumes that the observations and inputs are related according to a linear dynamical system:

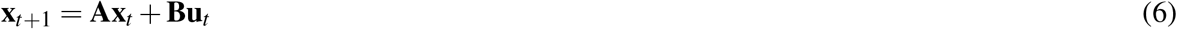

In general, when using LQR, the **A** and **B** matrices are known. However, our system dynamics are nonlinear, probabilistic, and change based on the neural encoder and decoder. Using naive system dynamics leads to a suboptimal feedback controller. To model system dynamics, we therefore used an algorithm to iteratively learn system dynamics **A** and **B** from the data collected while performing tasks. The data includes observation, **x**_*t*_, next observation, **x**_*t*+1_, and action, **u**_*t*_. We approximate the relationship between hand kinematics and cursor kinematics of BCI system with a linear dynamic system as:

##### Algorithm 1

Iteratively estimate system dynamics

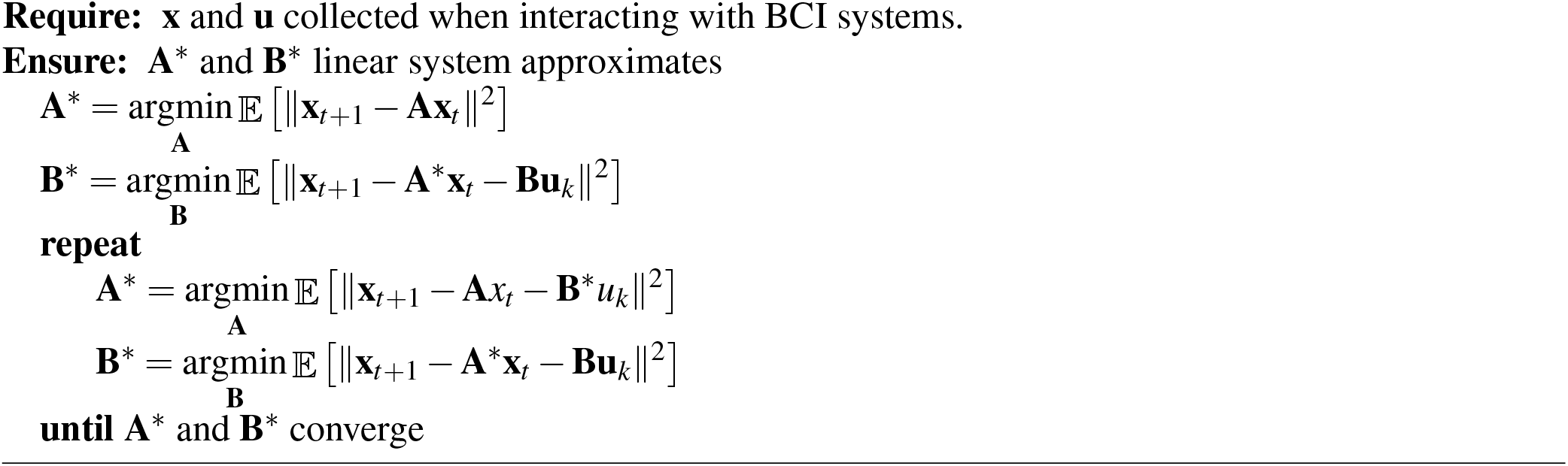

In our algorithm, we assume system state can evolve through its internal dynamics, **A**, without requiring external input. We therefore initialize **B** = 0. After inferring **A**^*∗*^ and **B**^*∗*^, we use the LQR algorithm to infer a linear policy, **u**_*t*_ = **Kx**_*t*_.

#### 4.7.2 Deep RL agent

To train the deep RL agent, we used proximal policy optimization (PPO) with new regularizations to constraint agent behavior as humans had. PPO uses a clipped surrogate objective function with a goal of reducing KL divergence between successive gradient updates^54^. To encourage exploration, the entropy of actions given state was added to the objective function. The objective function of PPO is defined as the following:

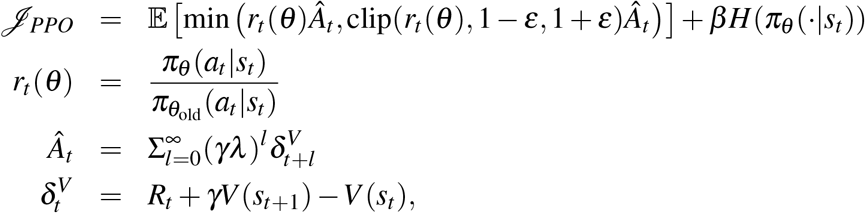

where *r*_*t*_ is the probability ratio, 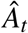 is the advantage value, *R*_*t*_ is the reward from environment, *V* (*s*_*t*_) is the output from the value function given *s*_*t*_, *γ* and *λ* are hyperparameters in generalized advantage estimation (GAE)^55^. PPO updates were performed with first-order stochastic gradient descent (SGD) or Adam. The agent outputs were the means and logarithmic standard deviations of Gaussian distributions modeling the hand acceleration. During testing, the actions injected into the tasks were sampled from Gaussian distributions on each dimension of action space.

The PPO algorithm included a policy and a value network. The policy network was a feedforward neural network with two affine layers of 64 neurons, using tanh(*·*) as the activation function, followed by a linear layer to output means and logarithmic standard deviation of Gaussian distributions modeling the 2D hand acceleration. The value network shared the same first two 64-neuron affine layers, followed by a linear layer to estimate the value function *V* (*s*_*t*_).

This PPO agent, naively trained, does not exhibit realistic physical behaviors, resulting in unmatched decoder performance (Supplementary Figure 11). For example, it could make arbitrarily large accelerations, something humans cannot do with their arms because of biomechanical constraints. We therefore developed two additional regularization terms to encourage the agent to act more like a BCI user would. We first incorporated a smoothness constraint that penalized the KL divergence on consecutive actions. Additionally, we incorporated a zeroness constraint that penalized changes to the policy that move policy away from a Gaussian distribution with zero mean and unit variance. This encouraged the network to conserve energy, not moving unless it was necessary. The objective function of our constrained PPO agent is therefore:

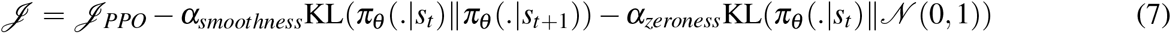

The policy network, *π*_*θ*_, outputs mean *µ* and logarithmic standard deviation, log *σ*, of a Gaussian distribution over the agent’s actions. We used the output of the policy network to estimate the KL divergence penalties. Consider two Gaussian distributions, *p*(*x*) and *q*(*x*), with means, *µ*_1_ and *µ*_2_, and standard deviations, *σ*_1_ and *σ*_2_, respectively. The KL divergence of these two Gaussian distributions is calculated in the following way:

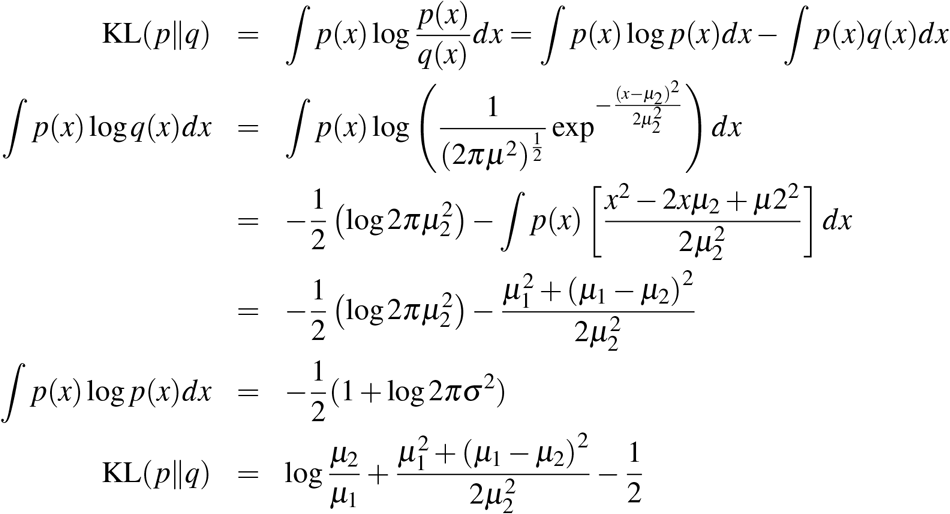

where, we used Var(*x*) = 𝔼 [*x*^2^] *−* 𝔼 [*x*]^2^ to replace ∫ *p*(*x*)*x*^2^ *dx* = *μ*^2^™*σ*^2^.

#### 4.7.3 Training RL agents with curriculum learning

We first trained an agent to perform the center-out-and-back task in a bypass mode which has cursor position equal to hand position. We adjusted the zero constraint and smooth constraint until the resulting behavior was close to Monkey J’s behavior. All other agents learned from this pretrained control policy. Although reward shaping facilitates the learning process, agents’ behaviors were also determined by the shaped reward. We therefore only trained agents with a simple reward setting which is 1 if agents successfully acquired targets and 0 others. We applied curriculum learning to facilitate the learning process by starting from a larger target acceptance window, 120 mm, and progressively shrunk it down to 30 mm if the overall target success rate was more than 90%.

## References

1. P R Kennedy and R A E Bakay. Restoration of neural output from a paralyzed patient by a direct brain connection. Neuroreport, 9(8):1707–1711, June 1998.

2. P R Kennedy, R A E Bakay, M M Moore, K Adams, and J Goldwaithe. Direct control of a computer from the human central nervous system. IEEE Trans. Rehabil. Eng., 8(2):198–202, June 2000.

3. Dawn M Taylor, Stephen I Helms Tillery, and Andrew B Schwartz. Direct cortical control of 3D neuroprosthetic devices. Science, 296(5574):1829–1832, June 2002.

4. Leigh R Hochberg, Mijail D Serruya, Gerhard M Friehs, Jon A Mukand, Maryam Saleh, Abraham H Caplan, Almut Branner, David Chen, Richard D Penn, and John P Donoghue. Neuronal ensemble control of prosthetic devices by a human with tetraplegia. Nature, 442(7099):164–171, July 2006.

5. Wei Wu, Yun Gao, Elie Bienenstock, John P Donoghue, and Michael J Black. Bayesian population decoding of motor cortical activity using a kalman filter. Neural Comput., 18(1):80–118, January 2006.

6. Leigh R Hochberg, Daniel Bacher, Beata Jarosiewicz, Nicolas Y Masse, John D Simeral, Joern Vogel, Sami Haddadin, Jie Liu, Sydney S Cash, Patrick van der Smagt, and John P Donoghue. Reach and grasp by people with tetraplegia using a neurally controlled robotic arm. Nature, 485(7398):372–375, May 2012.

7. Jennifer L Collinger, Brian Wodlinger, John E Downey, Wei Wang, Elizabeth C Tyler-Kabara, Douglas J Weber, Angus J C McMorland, Meel Velliste, Michael L Boninger, and Andrew B Schwartz. High-performance neuroprosthetic control by an individual with tetraplegia. Lancet, 381 (9866):557–564, February 2013.

8. Brian Wodlinger, J E Downey, Elizabeth C Tyler-Kabara, Andrew B Schwartz, M L Boninger, and Jennifer L Collinger. Ten-dimensional anthropomorphic arm control in a human brain-machine interface: difficulties, solutions, and limitations. J. Neural Eng., 12(1):016011, February 2015.

9. Vikash Gilja, Paul Nuyujukian, Cynthia A Chestek, John P Cunningham, Byron M Yu, Joline M Fan, Mark M Churchland, Matthew T Kaufman, Jonathan C Kao, Stephen I Ryu, and Krishna V Shenoy. A high-performance neural prosthesis enabled by control algorithm design. Nat. Neurosci., 15(12): 1752–1757, November 2012.

10. Joline M Fan, Paul Nuyujukian, Jonathan C Kao, Cynthia A Chestek, Stephen I Ryu, and Krishna V Shenoy. Intention estimation in brain-machine interfaces. J. Neural Eng., 11(1):016004, February 2014.

11. Maryam M Shanechi, Amy L Orsborn, Helene G Moorman, Suraj Gowda, Siddharth Dangi, and Jose M Carmena. Rapid control and feedback rates enhance neuroprosthetic control. Nat. Commun., 8:13825, 2017.

12. Jonathan C Kao, Paul Nuyujukian, Stephen I Ryu, Mark M Churchland, John P Cunningham, and Krishna V Shenoy. Single-trial dynamics of motor cortex and their applications to brain-machine interfaces. Nat. Commun., 6(May):1–12, 2015.

13. Jonathan C Kao, Paul Nuyujukian, Stephen I Ryu, and Krishna V Shenoy. A high-performance neural prosthesis incorporating discrete state selection with hidden markov models. IEEE Transactions on Biomedical Engineering, 64(4):935–945, April 2017.

14. Karunesh Ganguly and Jose M Carmena. Emergence of a stable cortical map for neuroprosthetic control. PLoS Biol., 7(7):e1000153, July 2009.

15. Amy L Orsborn, Helene G Moorman, Simon A Overduin, Maryam M Shanechi, Dragan F Dim-itrov, and Jose M Carmena. Closed-Loop decoder adaptation shapes neural plasticity for skillful neuroprosthetic control. Neuron, 82(6):1380–1393, June 2014.

16. Daniel B Silversmith, Reza Abiri, Nicholas F Hardy, Nikhilesh Natraj, Adelyn Tu-Chan, Edward F Chang, and Karunesh Ganguly. Plug-and-play control of a brain-computer interface through neural map stabilization. Nat. Biotechnol., 39(3):326–335, March 2021.

17. David Sussillo, Paul Nuyujukian, Joline M Fan, Jonathan C Kao, Sergey D Stavisky, Stephen I Ryu, and Krishna V Shenoy. A recurrent neural network for closed-loop intracortical brain-machine interface decoders. J. Neural Eng., 9(2):026027, April 2012.

18. David Sussillo, Sergey D Stavisky, Jonathan C Kao, Stephen I Ryu, and Krishna V Shenoy. Making brain–machine interfaces robust to future neural variability. Nat. Commun., 7:13749, December 2016.

19. Francis R Willett, Donald T Avansino, Leigh R Hochberg, Jaimie M Henderson, and Krishna V Shenoy. High-performance brain-to-text communication via handwriting, 2021.

20. Sung-Phil Kim, John D Simeral, Leigh R Hochberg, John P Donoghue, and Michael J Black. Neural control of computer cursor velocity by decoding motor cortical spiking activity in humans with tetraplegia. J. Neural Eng., 5(4):455–476, December 2008.

21. Steven M Chase, Andrew B Schwartz, and Robert E Kass. Bias, optimal linear estimation, and the differences between open-loop simulation and closed-loop performance of spiking-based brain-computer interface algorithms. Neural Netw., 22(9):1203–1213, 2009.

22. Shinsuke Koyama, Steven M Chase, Andrew S Whitford, Meel Velliste, Andrew B Schwartz, and Robert E Kass. Comparison of brain-computer interface decoding algorithms in open-loop and closed-loop control. J. Comput. Neurosci., 29(1-2):73–87, August 2010.

23. John P Cunningham, Paul Nuyujukian, Vikash Gilja, Cindy A Chestek, Stephen I Ryu, and Krishna V Shenoy. A closed-loop human simulator for investigating the role of feedback control in brain-machine interfaces. J. Neurophysiol., 105(4):1932–1949, April 2011.

24. Sergey D Stavisky, Jonathan C Kao, Paul Nuyujukian, Stephen I Ryu, and Krishna V Shenoy. A high performing brain–machine interface driven by low-frequency local field potentials alone and together with spikes. J. Neural Eng., 12(3):036009, 2015.

25. Francis R Willett, Daniel R Young, Brian A Murphy, William D Memberg, Christine H Blabe, Chethan Pandarinath, Sergey D Stavisky, Paymon Rezaii, Jad Saab, Benjamin L Walter, Jennifer A Sweet, Jonathan P Miller, Jaimie M Henderson, Krishna V Shenoy, John D Simeral, Beata Jarosiewicz, Leigh R Hochberg, Robert F Kirsch, and A Bolu Ajiboye. Principled BCI decoder design and parameter selection using a feedback control model. Sci. Rep., 9(1):8881, June 2019.

26. Manuel Lagang and Lakshminarayan Srinivasan. Stochastic optimal control as a theory of brain-machine interface operation. Neural Comput., 25(2):374–417, February 2013.

27. S Gowda, A L Orsborn, S A Overduin, H G Moorman, and J M Carmena. Designing dynamical properties of Brain–Machine interfaces to optimize Task-Specific performance. IEEE Trans. Neural Syst. Rehabil. Eng., 22(5):911–920, September 2014.

28. Maryam M Shanechi, Ziv M Williams, Gregory W Wornell, Rollin C Hu, Marissa Powers, and Emery N Brown. A real-time brain-machine interface combining motor target and trajectory intent using an optimal feedback control design. PLoS One, 8(4):23–32, 2013.

29. Maryam M Shanechi, Amy L Orsborn, and Jose M Carmena. Robust brain-machine interface design using optimal feedback control modeling and adaptive point process filtering. PLoS Comput. Biol., 12 (4):e1004730, 2016.

30. Miri Benyamini and Miriam Zacksenhouse. Optimal feedback control successfully explains changes in neural modulations during experiments with brain-machine interfaces, 2015.

31. Yin Zhang and Steve M Chase. Optimizing the usability of Brain-Computer interfaces. Neural Comput., 30(5):1323–1358, May 2018.

32. Francis R Willett, Chethan Pandarinath, Beata Jarosiewicz, Brian A Murphy, William D Memberg, Christine H Blabe, Jad Saab, Benjamin L Walter, Jennifer A Sweet, Jonathan P Miller, Jaimie M Henderson, Krishna V Shenoy, John D Simeral, Leigh R Hochberg, Robert F Kirsch, and A Bolu Aji-boye. Feedback control policies employed by people using intracortical brain–computer interfaces. J. Neural Eng., 14(1):016001, November 2016.

33. Ken-Fu Liang and Jonathan C Kao. Deep learning neural encoders for motor cortex. IEEE Trans. Biomed. Eng., 67(8):2145–2158, August 2020.

34. Siddharth Dangi, Amy L Orsborn, Helene G Moorman, and Jose M Carmena. Design and analysis of closed-loop decoder adaptation algorithms for brain-machine interfaces. Neural Comput., 25(7): 1693–1731, July 2013.

35. Byron M Yu, John P Cunningham, Gopal Santhanam, Stephen I Ryu, and Krishna V Shenoy. Gaussian-process factor analysis for low-dimensional single-trial analysis of neural population activity. J. Neurophysiol., 102:612–635, 2009.

36. John P Cunningham and Byron M Yu. Dimensionality reduction for large-scale neural recordings. Nat. Neurosci., 17(11):1500–1509, August 2014.

37. Juan A Gallego, Matthew G Perich, Lee E Miller, and Sara A Solla. Neural manifolds for the control of movement. Neuron, 94(5):978–984, June 2017.

38. Mark M Churchland, John P Cunningham, Matthew T Kaufman, Justin D Foster, Paul Nuyujukian, Stephen I Ryu, and Krishna V Shenoy. Neural population dynamics during reaching. Nature, 487 (7405):51–56, July 2012.

39. Chethan Pandarinath, Daniel J O’Shea, Jasmine Collins, Rafal Jozefowicz, Sergey D Stavisky, Jonathan C Kao, Eric M Trautmann, Matthew T Kaufman, Stephen I Ryu, Leigh R Hochberg, Jaimie M Henderson, Krishna V Shenoy, L F Abbott, and David Sussillo. Inferring single-trial neural population dynamics using sequential auto-encoders, 2018.

40. Guillaume Hennequin, Tim P Vogels, and Wulfram Gerstner. Optimal control of transient dynamics in balanced networks supports generation of complex movements. Neuron, 82(6):1394–1406, June 2014.

41. David Sussillo, Mark M Churchland, Matthew T Kaufman, and Krishna V Shenoy. A neural network that finds a naturalistic solution for the production of muscle activity. Nat. Neurosci., 18(7):1025–1033, 2015.

42. Jonathan A Michaels, Benjamin Dann, and Hansjörg Scherberger. Neural population dynamics during reaching are better explained by a dynamical system than representational tuning. PLoS Comput. Biol., 12(11):e1005175, 2016.

43. Jonathan C Kao. Considerations in using recurrent neural networks to probe neural dynamics. J. Neurophysiol., 122(6):2504–2521, December 2019.

44. Florian Fischer, Miroslav Bachinski, Markus Klar, Arthur Fleig, and Jörg Müller. Reinforcement learning control of a biomechanical model of the upper extremity. Sci. Rep., 11(1):14445, July 2021.

45. Emanuel Joos, Fabien Péan, and Orcun Goksel. Reinforcement learning of musculoskeletal control from functional simulations. July 2020.

46. Jan Peters and Stefan Schaal. Reinforcement learning of motor skills with policy gradients. Neural Netw., 21(4):682–697, May 2008.

47. Seungmoon Song, Lukasz Kidzinski, Xue Bin Peng, Carmichael Ong, Jennifer Hicks, Sergey Levine, Christopher G Atkeson, and Scott L Delp. Deep reinforcement learning for modeling human locomo-tion control in neuromechanical simulation. August 2020.

48. Aravind Rajeswaran, Vikash Kumar, Abhishek Gupta, Giulia Vezzani, John Schulman, Emanuel Todorov, and Sergey Levine. Learning complex dexterous manipulation with deep reinforcement learning and demonstrations. September 2017.

49. Alan D Degenhart, William E Bishop, Emily R Oby, Elizabeth C Tyler-Kabara, Steven M Chase, Aaron P Batista, and Byron M Yu. Stabilization of a brain–computer interface via the alignment of low-dimensional spaces of neural activity. Nature Biomedical Engineering, April 2020.

50. Noshaba Cheema, Laura A Frey-Law, Kourosh Naderi, Jaakko Lehtinen, Philipp Slusallek, and Perttu Hämäläinen. Predicting Mid-Air interaction movements and fatigue using deep reinforcement learning. In Proceedings of the 2020 CHI Conference on Human Factors in Computing Systems, CHI ‘20, pages 1–13, New York, NY, USA, April 2020. Association for Computing Machinery.

51. Jonathan C Kao, Sergey D Stavisky, David Sussillo, Paul Nuyujukian, and Krishna V Shenoy. Information systems opportunities in brain-machine interface decoders. Proc. IEEE, 102(5):666–682, 2014.

52. David Sussillo and Larry F Abbott. Generating coherent patterns of activity from chaotic neural networks. Neuron, 63(4):544–557, 2009.

53. Pavan Mehrotra, Sabar Dasgupta, Samantha Robertson, and Paul Nuyujukian. An open-source realtime computational platform (short WIP paper). ACM SIGPLAN Notices, 53(6):109–112, 2018.

54. John Schulman, Filip Wolski, Prafulla Dhariwal, Alec Radford, and Oleg Klimov. Proximal policy optimization algorithms. July 2017.

55. John Schulman, Philipp Moritz, Sergey Levine, Michael Jordan, and Pieter Abbeel. High-dimensional continuous control using generalized advantage estimation, 2015. URL https://arxiv.org/abs/1506.02438.

